# Shear stress and very low levels of ligand synergize to activate ALK1 signaling in endothelial cells

**DOI:** 10.1101/2023.11.01.565194

**Authors:** Ya-Wen Cheng, Anthony R. Anzell, Tristin A. Schwartze, Cynthia S. Hinck, Andrew P. Hinck, Beth L. Roman, Lance A. Davidson

**Affiliations:** Department of Bioengineering, Swanson School of Engineering, University of Pittsburgh, Pittsburgh, PA 15261, USA; Department of Human Genetics, School of Public Health, University of Pittsburgh, Pittsburgh, PA 15261, USA; Department of Structural Biology, School of Medicine, University of Pittsburgh, Pittsburgh, PA 15261, USA; Heart, Lung, Blood and Vascular Medicine Institute, University of Pittsburgh, Pittsburgh, PA 15261, USA; Department of Developmental Biology, School of Medicine, University of Pittsburgh, Pittsburgh, PA 15261, USA; Department of Computational and Systems Biology, School of Medicine, University of Pittsburgh, Pittsburgh, PA 15261, USA

**Keywords:** endothelial cell biology, mechanotransduction, mechanobiology, mechanosensing, pSMAD1/5/9, ALK1 receptor trafficking, hereditary hemorrhagic, telangiectasia

## Abstract

Endothelial cells (ECs) respond to concurrent stimulation by biochemical fac-tors and wall shear stress (SS) exerted by blood flow. Disruptions in flow-induced responses can result in remodeling issues and cardiovascular diseases, but the detailed mechanisms linking flow-mechanical cues and biochemical signaling remain unclear. Activin receptor-like kinase 1 (ALK1) integrates SS and ALK1-ligand cues in ECs; ALK1 mutations cause hereditary hemorrhagic telangiectasia (HHT), marked by arteriovenous malformation (AVM) development. However, the mechanistic underpinnings of ALK1 signaling modulation by fluid flow and the link to AVMs remain uncertain. We recorded EC responses under varying SS magnitudes and ALK1 ligand concentrations by assaying pSMAD1/5/9 nuclear localization using a custom multi-SS microfluidic device and a custom image analysis pipeline. We extended the previously reported syn-ergy between SS and BMP9, to include BMP10 and BMP9/10 . Moreover, we demonstrated this synergy is effective even at extremely low SS magnitudes (0.4 dyn/cm2) and ALK1 ligand range (femtogram/mL). The synergistic response to ALK1 ligands and SS requires the kinase activity of ALK1. Moreover, ALK1’s basal activity and response to minimal ligand levels depend on endo-cytosis, distinct from cell-cell junctions, cytoskeleton-mediated mechanosensing, or cholesterol-enriched microdomains. Yet, an in-depth comprehension of ALK1 receptor trafficking’s molecular mechanisms requires further investigation.

## Introduction

Endothelial cells (EC) that line blood vessels can sense the wall shear stress (SS) exerted by blood flow [1–3]. EC responses to these fluidic mechanical cues include dilation or contraction of vessels [4, 5] as well as angiogenesis and vascular remodeling [6–8]. Whereas pulsatile laminar flow induces quiescence and is atheroprotective, oscillatory or disturbed flow can lead to cardiovascular diseases such as atherosclerosis [9]. Although numerous studies have been carried out at the interface between EC biology and biophysics, we still do not understand the key features of the EC response to flow, or how ECs integrate flow-mediated mechanical cues with the variety of biochemical cues within their microenvironment.

One way in which mechanical cues are thought to regulate EC biology is by modifying the sensitivity of biochemical signaling pathways. Previous studies had identified a role for vascular endothelial growth factor receptors (VEGFRs) in the EC response to flow. Under static conditions, VEGFA binding to a VEGFR2 dimer causes the tyrosine kinase in the dimer complex to be transactivated; activated VEGFR2 can then drive Src, ERK, and PI3K signaling cascades [10]. When blood vascular ECs expressing platelet endothelial cell adhesion molecule (PECAM, or CD31) and vascular endothelial cadherin (VE-cadherin, or CDH5) are subjected to flow, VEGFR2 becomes phosphorylated and can activate PI3K even in the apparent absence of VEGFA ligand [11, 12], leading to EC elongation along the flow axis and Golgi-nuclear polarization and orientation against the direction flow [13]. A related receptor, VEGFR3, is expressed at relatively high levels in lymphatic ECs compared to blood ECs, the former of which experience comparatively low SS. Like VEGFR2, VEGFR3 can be activated by SS in a ligand-independent manner, and it is necessary and sufficient for EC orientation with flow at low SS magnitudes (∼5 dyn/cm2) [14, 15]. Based on these studies, it has been proposed that ECs have a flow preference, i.e. . a “set point” or preferred magnitude of SS, which is inversely correlated with the protein level of VEGFR3 [15]. At the preferred SS, ECs align along the flow axis and remain stable and quiescent; above or below this SS level, ECs mount an inflammatory response and initiate vascular remodeling to bring SS back to the desired set point [16].

SS stimulation is also thought to enhance the ligand-dependent activity of activin receptor-like kinase 1 (ALK1). ALK1 is one of seven TGF𝛽 family type I receptor serine/threonine kinases; it is the only type I receptor that strongly binds circulating ligands bone morphogenetic protein 9 (BMP9) and BMP10. Ligand binding brings together a heterotetrameric complex at the plasma membrane containing two ALK1 and two type II receptors, allowing the type II receptors to phosphorylate and activate ALK1. ALK1 then phosphorylates receptor-specific (r-) SMADs, SMAD1, SMAD5, and/or SMAD9 (hereafter, SMAD1/5/9). Two phosphorylated r-SMAD molecules complex with one molecule of the common partner, SMAD4, and the complex subsequently undergoes nuclear localization and acts as a transcription factor [17–21]. Type I receptors ALK2, ALK3, and ALK6 bind alternative BMPs but similar to ALK1, phosphorylate SMAD1/5/9 [22].

ALK1 function is required in vivo and in vitro for ECs to orient optimally against flow, i.e. for the vector from the nucleus to the Golgi to be directed upstream, against the flow vector. ALK1 is also required for flow-induced upstream migration of mouse retinal capillary and venous ECs [23]. Moreover, alk1 expression in zebrafish arterial ECs depends on flow, and ECs lacking alk1 fail to migrate against flow and exhibit dysregulated expression of several flow-sensitive genes [24, 25]. Thus, like VEGFRs, ALK1 appears to play an important role in integration of mechanical and biochemical cues that shape EC flow response pathways.

Defects in ALK1 signaling are thought to lead to vascular disease by disrupting EC responses to SS at different time scales. Under normal conditions, ALK1 signaling impacts EC behavior on both short and long time scales; pSMAD1/5/9 nuclear localization and downstream gene expression on timescales of less than an hour [26] and EC migration during development over hours to days [25]. Defects in ALK1 signaling, perhaps as a consequence of defects in these processes, lead to arteries and veins pathologically forming direct connections known as arteriovenous malformations (AVMs), which decrease gas exchange and may be prone to hemorrhage [27]. Such AVMs are common features of hereditary hemorrhagic telangiectasia (HHT), an autosomal dominant vascular disorder that impacts approximately 1 in 5000 people worldwide and is correlated in more than 90% of cases to mutations in ALK1 or ENG (encoding endoglin, a non-signaling BMP9/BMP10 receptor) [28]. Additionally, ALK1 signaling deficiencies are associated with rare cases of pulmonary arterial hypertension [29, 30]. Mutations in SMAD4 can also cause HHT [31], and SMAD4, is required for ECs to exhibit the features of flow-mediated morphological responses within their set point range of fluid SS [16].

Given the challenges in controlling in vivo flow conditions, efforts to study EC responses to SS have driven the development of in vitro flow models. Parallel plate flow chambers [11, 15, 26, 32] and orbital shakers [33, 34] are most commonly used to deliver flow cues. While orbital shakers generate a range of fluid flow magnitudes and directions, parallel plate flow devices can deliver precise levels of SS. However, a major practical limitation of parallel plate systems is that they typically require more than 200 mL of medium per experiment, which may preclude their use with rare or costly reagents such as recombinant growth factors. Thus, efforts to study the combined impact of flow and biochemical cues have turned to microphysiological systems based on microfluidic devices. Microfluidic devices can generate multiple flow regimes and limit the amount of media needed per run [32, 35–37]. Furthermore, if designed for operation on the stage of a compound microscope, microfluidic devices can allow real-time or post-experimental imaging and high-throughput quantification at the single cell level. In contrast to bulk methods such as

Western blot, imaging can provide extensive details on cell-to-cell variation and how cells integrate signaling from SS and growth factors [11, 15, 26].

In most microphysiological studies to date, SS regimes have mainly focused on the physiological range of arterial flow, 10 – 50 dyn/cm2 [38, 39]; however, it is also critical to understand how ECs respond to flows outside this range, which exist in non-arterial vessels, in vascular pathologies, and during vessel remodeling [40]. Yet, few studies have examined EC responses to low laminar SS (< 5 dyn/cm2) found in veins or lymphatic vessels [13, 15, 41–44]. Consequently, we have a limited understanding of EC biology at the lowest level of SS that ECs can sense.

In this paper, we apply a custom multi-SS microfluidic device [37] combined with an image analysis pipeline to quantify EC responses to ALK1-ligands under flow and to test mechanisms responsible for synergistic interactions between biomechanical flow and biochemical stimuli. We recorded EC responses at single-cell resolution by measuring pSMAD1/5/9 nuclear localization. Our analysis confirms the previously observed synergy between SS and BMP9 and extends these findings to ALK1 ligands, BMP10 and BMP9/10. Additionally, we demonstrate that synergy occurs at very low SS magnitudes, as low as 0.4 dyn/cm^2^, and with ultra-low levels of ALK1 ligand, estimated in the femtogram/mL range. The synergetic response to ALK1 ligands and SS requires the kinase activity of the ALK1 receptor but not cell-cell adhesion, and it is refractory to modest perturbation of the actin cytoskeleton or cholesterol-enriched microdomains on cell membranes. However, basal activity of ALK1 and its response to ultra-low ligand depends on endocytosis.

## Materials and methods

### Cell culture

Pooled human umbilical vein ECs (HUVECs; Cat: C-12203, Promocell, Heidelberg, Germany) or human pulmonary artery ECs (HPAECs; Cat: C-12241, Promocell) were cultured in a sterile humidified incubator in complete endothelial cell growth medium (EC Growth Medium 2/EGM2, containing 2% FBS; Cat: C-22011, Promocell) and 1 × antibiotic-antimycotic (Cat: 15240112, Thermo Fisher Scientific, Waltham, MA USA) at 37 ° C under 5% CO2. Upon confluency, cells were rinsed with HEPES BSS (Detach Kit, Cat: C-41220, Promocell) and treated with 0.04% trypsin/0.03% EDTA for 5-7 minutes at room temperature until cells detached. After adding trypsin neutralization solution, cells were centrifuged at 220 × g for 3 minutes and gently resuspended in fresh EGM2. Cells were maintained in EGM2 for a maximum of six passages.

### shRNA transduction

Lentiviral particles containing shRNA targeting ALK1 exon 11 (shALK1, Cat: TRCN0000000355; tccagagaagcctaaagtgat) or non-targeted control (shCont, Cat: SHC002) were generated in HEK293 cells using the Sigma Mission system (Sigma-Aldrich, St. Louis, MO USA). HUVECs were seeded into a 6-well dish at a density of 80,000 cells per well. The next day (∼50% confluency), cells were transduced with 30 MOI lentivirus, selected with 1 µg/mL puromycin for 5 days before seeding into devices, and maintained in 1 µg/mL puromycin throughout the experiment. To determine knockdown (KD) efficiency, RNA and protein were extracted 6 days post-transduction using the AllPrep RNA/Protein kit (Cat: 80204, Qiagen, Germantown, MD USA). RNA was converted to cDNA using SuperScript IV cDNA Synthesis Kit (Cat: 18091050, Invitrogen/Thermo Fisher Scientific) and analyzed by RT-qPCR on a Quant Studio 12K Flex Real Time PCR system, version 1.1.2 (Applied Biosystems/Life Technologies/Thermo Fisher Scientific) using validated (efficiency > 1.8) TaqMan assays (Thermo Fisher Scientific) for ALK1 (Cat: Hs00953798_m1) and b-ACTIN (Cat: Hs0106066575_g1), with data analyzed via the DDCt method [45]. For protein analysis, lysates were analyzed using the Pierce BCA Protein Assay Kit (Cat: 23225; Thermo Fisher Scientific). Ten µg of protein was separated by reducing SDS-PAGE, transferred to nitrocellulose, and probed with antibodies targeting ALK1 [Rabbit anti-human ALK1 [46], 1:1,000, 4°C overnight; gift provided by D. Marchuk, Duke University] and GAPDH (mouse anti-human GAPDH, 1:10,000, 1 hr at RT; Cat: ab8245, Abcam, Waltham, MA). Primary antibodies were detected using donkey anti-rabbit IRDye® 800CW IgG (Cat: 925-32213) and goat anti-mouse IgG IRDye® 680LT (Cat: 926-68020), respectively (LI-COR, Lincoln, NE USA) and blots imaged using a LI-COR Odyssey Clx infrared imaging system and analyzed using Image Studio v5.2.

### EC seeding and SS experiments

Cell seeding and SS experiments were similar to our previous study [37] and are briefly summarized below. To ensure a confluent monolayer, ECs were seeded at a density of 3 million/mL in a volume of 40 µl in a fibronectin-coated microfluidic device, which was placed in a culture plate. For isolated cell experiments, ECs were seeded at a density of 0.5 million/mL. To prevent drying out, an additional 0.5 mL EGM2 was added to the top of the inlet and outlet ports of the device. Subsequently, devices were placed in the culture plates, and the lids were closed. The cells were cultured for at least 16 hours.

After 16 hours of growth, all cultures were serum starved in 0.2% BSA medium [30% BSA (Cat: A9576, Sigma-Aldrich) diluted in EC Basal Medium-2 (EBM2), phenol red-free (Cat: C-22216, Promocell)] for 4 hours. For flow conditions, cultures were subsequently subjected to SS with a circulating flow (∼13 mL reservoir per channel) driven by an Ismatec® Reglo independent channel control 4-channel peristaltic pump (Cat: ISM4408, Cole-Parmer, Vernon Hills, IL USA) for 45 minutes. For the dose response experiments, BMP9 (Cat: 3209-BP, R&D Systems, Minneapolis, MN USA), BMP10 (Cat: 2926-BP, R&D Systems), and BMP9/10 heterodimer (see below for preparation) were diluted in 0.2% BSA medium. Flow direction is right to left for all of the images.

For static conditions within the device, a medium exchange was carried out using 0.2% BSA medium with the addition of BMP (experimental) or without (control). This exchange was accomplished by pipetting 0.2 mL of the medium into the inlet and outlet ports, and0.5 mL of same medium to the top of the device, effectively covering the inlet and outlet ports to prevent drying out.

For static conditions within the plate, we performed static experiments in four-compartment glass bottom culture plates (Cat: 627871, Greiner Bio-One, Monroe, NC, USA or Cat: D35C4-20-1.5-N, Cellvis, Mountain View, CA, USA, if not specified). Cells were grown for 16 hrs followed by 4 hr starvation before treatment.

At the end of each experiment, frozen 4% paraformaldehyde (PFA) was freshly thawed at 37°C until fully dissolved. Cells were fixed with 4% PFA for 15 minutes and stored in PBS at 4°C until immunofluorescence staining within less than 5 days.

### BMP9/10 heterodimer purification

cDNA encoding human full length pro-BMP9 (NP_057288.1, amino acids 23-429) or pro- BMP10 (NP_055297.1, amino acids 22-424) was cloned into pcDNA3.1+ (Invitrogen/Thermo Fisher Scientific) downstream of the rat serum albumin signal peptide, a hexahistidine tag (HHHHHH; BMP9) or strepII tag (WSHPQFEK; BMP10), and a factor Xa cleavage site (IEGR). To prevent disassociation during purification and ensure complete processing, the natural primary furin processing site separating the pro and growth factor domains were replaced with a factor Xa processing site (IEGR in place of RRKR in pro-BMP9, and GIEGR in place of AIRIRR in pro- BMP10). Coding sequences were obtained by gene synthesis (Twist Biosciences, South San Francisco, CA USA) and inserted between either the NheI and XhoI or NheI and ApaI sites in pcDNA3.1+. Constructs were verified over their entire length by sequencing in the forward and reverse directions using primers that bound to the CMV promoter or the BGH terminator.

The plasmids coding for pro-BMP9 and pro-BMP10 were co-transected in equal amounts into HEK293 cells (expi293, Invitrogen/Thermo Fisher Scientific) and after five days, the conditioned medium was harvested by removing the cells by centrifugation as previously described [47]. The medium was dialyzed into IMAC column buffer (25 mM sodium phosphate, 150 mM NaCl, 8 mM imidazole, pH 8.0) before being loaded onto a nickel-loaded chelating sepharose column (Cytiva, Marlborough, MA USA). The protein was eluted over a 0% to 100% (0.5 M imidazole) gradient over 300 mL, and the fractions containing recombinant pro-BMP9/10 dimers and monomer were identified by non-reducing SDS-PAGE and pooled. The sample was dialyzed into 100 mM Tris, 150 mM NaCl, 1 mM EDTA, pH 8.0, loaded onto a streptactin II column (IBA Life Sciences, Germany), washed with 10 column volumes to remove unbound protein, and eluted with 50 mM D-biotin.

The eluate was dialyzed into 20 mM Tris pH 8.0, 100 mM NaCl, 2 mM CaCl2, 0.02% NaN3 and digested with factor Xa at 37 °C at a 1000:1 target protein:factor Xa ratio over two days. The digest mixture was dialyzed into 4 mM HCl before being loaded onto a 10 x 250 mm 2 μm C18 Jupiter reverse phase column (Phenomenex, Torrance, CA USA) equilibrated in 95% A/5% B (where A and B are water and acetonitrile with 0.1% TFA). The monomers and dimers were eluted separately over a gradient from 30% to 40% B over 400 mL. The resulting fractions were then lyophilized and resuspended in 50 mM acetic acid.

### Small molecule inhibitors and ALK1-depletion

For inhibitor or drug test, most chemicals were reconstituted in DMSO and diluted with 0.2% BSA medium. However, MβCD was directly dissolved in the 0.2% BSA medium. These samples were then applied during the serum starvation period, prior to the application of flow. The following concentrations and incubation times were utilized for each chemical: LDN193189 (1 µM; Cat: S2618, Selleck Chemicals, Houston, TX USA) and Y27632 (10 µM; Cat: 688000, Calbiochem, San Diego, CA USA) were applied for 60 minutes prior to flow. Latrunculin B (0.015 µM; Cat: 428020, Calbiochem), MβCD (5 mM; Cat: 377110050, Thermo Fisher Scientific) and Dynasore (100 µM; Cat: D7693, Sigma-Aldrich) were applied for 30 minutes prior to flow. Controls were treated with the same percentage of DMSO in 0.2% BSA medium as the inhibitor or drug-treated samples: LDN193189 control, 0.2% DMSO; Y27632 control, 0.1% DMSO; Latrunculin B and Dynasore controls, 0.4% DMSO; MβCD control, 0% DMSO. ALK1-Fc (Cat: 370-AL, R&D Systems) was incubated at a final concentration of 1.25 mg/mL with 30% BSA on an orbital shaker at 4°C overnight. ALK1-Fc-complexes were then captured with CaptureSelect FcXL Affinity Matrix (Cat: 194328010, Thermo Fisher Scientific). The ALK1 ligand-depleted 30% BSA was diluted to 0.2% with phenol red-free EBM2 for use.

### Immunofluorescence

For immunofluorescence, all steps were performed within the microfluidic device at room temperature unless otherwise stated. Cells were permeabilized with 0.1% (v/v) Triton X-100 (Cat: T8787, Sigma-Aldrich) in PBS (PBST) for 15 minutes and incubated with 10% (v/v) goat serum (Cat: G9023, Sigma Aldrich) in PBST for 30 minutes. Cells were incubated overnight at 4° C with rabbit anti-pSMAD1/5/9 (1:600 dilution, Cat: 13820, Cell Signaling Technology, Danvers, MA USA). The secondary antibodies: goat anti-rabbit IgG-FITC (Cat: 111-095-045) or goat anti-rabbit IgG-Rhodamine Red™-X (Cat: 111-295-045) (Jackson ImmunoResearch Laboratories, Westgrove, PA USA) were diluted 1:500 in PBST and applied for 1 hour. Cells were then washed with PBST and simultaneously stained for 10 minutes for nuclei (DAPI) and, when needed, actin filaments (Acti-stain 670 phalloidin, 1:600 dilution; Cat: SKU PHDN1, Cytoskeleton, Denver, CO USA). Imaging was performed using a spinning disk confocal microscope (Yokogawa CSU-X1, Hamamatsu X2 EMCCD mounted on a Leica DMI-6000).

### Image analysis of ALK1 signaling

Images of four fields were analyzed per region of each device or plate, which included approximately 80-120 cells per run. To quantify ALK signaling, we modified a custom-written macro for Fiji [48] that we developed previously [37]. This macro was altered to quantify relative amounts of pSMAD1/5/9 in nuclei and nuclei-adjacent cytoplasm. In brief, the DAPI channel was used to segment regions of interest (ROI) containing individual nuclei. Rather than use another channel to segment cell boundaries, we simplified the analysis by measuring pSMAD1/5/9 signal in cytoplasm surrounding the nucleus. This strategy limited measurement of pSMAD1/5/9 in regions where ECs overlap. Thus, to quantify cytosolic pSMAD1/5/9, we generated an annular region surrounding the nucleus in each cell from the nuclear ROI. The annulus consisted of a three-micron-wide band outside each nucleus ROI. To test whether this annulus included high levels of peri- or juxta-nuclear pSMAD1/5/9, we varied the distance of the inner ring of the annular band from the nuclear ROI and found no or minimal significant difference over moderate distances. The ALK signaling index (ASI) represents the level of pSMAD1/5/9 nuclear localization activated by any BMP type 1 receptor. We measured pSMAD1/5/9 median intensities within the nuclear (pSMADNuc) and cytosolic (pSMADCyt) ROIs in each cell (Fig. 1B) and calculated:

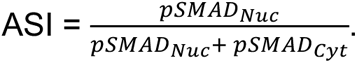

**Figure 1.**
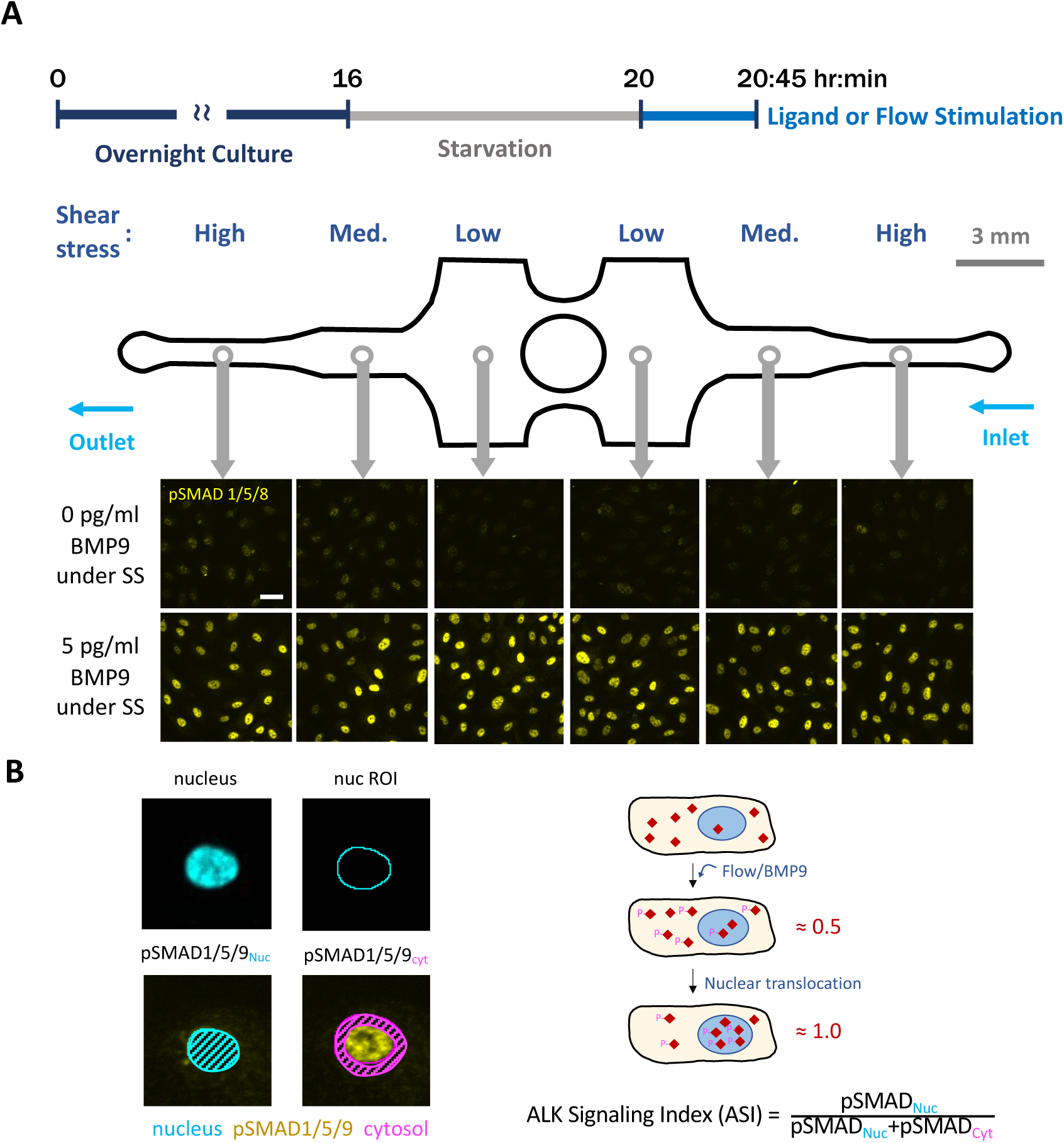
Experimental timeline, microfluidic device & image analysis pipeline. (A) General experiment design: Cells were cultured in complete growth medium under static conditions for ∼ 16 hours until confluency, starved for 4 hours with 0.2% BSA medium, and stimulated with or without serum/ligands and flow for 45 mins. Based on their position in the device, cells were exposed to high, medium, or low SS, as determined by the channel width. pSMAD1/5/9 immunofluorescence was used as a proxy for ALK1 signaling. Scale bar = 40 µm. (B) The ALK1 signaling index (ASI) of each cell was calculated using a customized image analysis pipeline as the ratio of pSMAD1/5/9 nuclear median intensity to pSMAD1/5/9 nuclear + cytoplasmic median intensity. Cytoplasmic fluorescence intensity was measured within a 3-µm annulus surrounding the nucleus.

This analysis is based on the assumption that these SMADs are distributed homogeneously in the cytosol and nucleus at dynamic equilibrium, returning an ASI of 0.5. Once ligand or SS is applied, SMAD1/5/9 will be phosphorylated, translocated into the nucleus, and increase the ASI. When the signaling reaches saturation, most pSMAD1/5/9 will be in the nucleus, with the ASI close to 1.0. Therefore, the image analysis pipeline allows an automated analysis of cellular response in terms of ALK signaling.

### Validation of disruption of actomyosin cell contractility

To quantify the morphology of cell–cell junctions after Rho Kinase (ROCK) inhibition by Y27632, we measured the tortuosity of cell boundaries [49]. The F-actin cytoskeleton was first labeled with phalloidin (see Immunofluorescence). Each cell in the confluent EC monolayer contacts immediately adjacent cells along bicellular and tricellular junctions, i.e. vertices. To quantify morphology, we measured linear distance between vertices (P) and the pathlength along bicellular junctions (L). The tortuosity index then was calculated by the following equation:

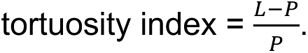

### Validation of F-actin level reduction

To quantify the effectiveness of latrunculin B in reducing F-actin levels [50], confluent HUVECs were serum starved for 4 hours in 0.2% BSA medium, then treated with specified concentrations of latrunculin B for 30 minutes in 0.2% BSA medium. Subsequently, cells were treated with the same latrunculin B concentrations along with 1 pg/mL BMP9 for 45 minutes, fixed, and stained with phalloidin (see Immunofluorescence). For F-actin quantification, two Z-planes (2 µm interval) closest to apical and basal sides were analyzed for actin intensity respectively. Three images from each condition for each experiment were captured using a spinning disk confocal microscope (as described above). Actin intensity was Z-projected by average intensity and quantified as the median across all fields using Fiji software.

### Validation of lipid raft disruption

To quantify the effectiveness of lipid raft disruption by cholesterol chelation, we visualized raft lipids with Cholera toxin subunit B (CtxB) in HUVECs [51, 52]. First, HUVECs were seeded onto Millicell EZ 4-well glass chamber slides (Cat: PEZGS0816, Millipore Sigma) at a density of 80,000 cells per well after pre-coating with 25 µg/mL fibronectin (Cat: F1141, Sigma Aldrich). Cells were treated with specified concentrations of MβCD for 75 minutes in 0.2% BSA medium, washed with PBS, fixed with 4% PFA, and stained with CtxB, Alexa Fluor 647 conjugate (Cat: C34778, Invitrogen/Thermo Fisher Scientific, 1:30 dilution). Cells were then washed with PBST and mounted with Vectashield anti-fade mounting medium with DAPI (Cat: H-1800, Vector Laboratories). Nine images from each condition for each experiment were captured using a spinning disk confocal microscope (as described above). Images were analyzed in Fiji using a custom macro to count the number of CtxB puncta, which was normalized to the number of nuclei. In brief, max intensity z-projection was performed for each image followed by segmentation of CtxB puncta via autothreshold (set to “default”) and “watershed” separation to isolate overlapping puncta. Puncta were then counted using the “Analyze particles” command with a size exclusion criteria of anything < 1 µm2.

### Statistical analysis

Each experiment was repeated at least three independent times and the image analysis pipeline calculated the quantitative parameters for each experiment independently. Ligand dose response curves were generated and EC50 values were calculated by nonlinear sigmoidal 4PL fitting; EC50 values were compared across treatments by extra sum-of-squares F test [53]. Data from two different treatments were compared with a two-tailed unpaired student’s t-test. Experiments with a single variable, such as SS magnitude or the distance between nuclear and cytosol ROI, but more than one condition were analyzed by one-way ANOVA and Tukey multiple comparisons test. Drug and ALK1 KD experiments were analyzed using two-way ANOVA and Sidak’s or Tukey multiple comparisons test. Experiments with two variables, such as static versus SS and vehicle versus drug-treated, were analyzed by two-way ANOVA.

## Results

### Experimental design and image analysis pipeline

To understand how SS interacts with ALK1 signaling, we used a multimodal microfluidic device that we developed previously to deliver precise SS cues to ECs [37]. Our device integrates three physiological uniform laminar SS regions into a single chamber with varying channel widths and a uniform channel height (Fig. 1A). We seeded ECs into devices and cultured cells until confluency under static, no-flow conditions. We then serum starved cultures and subsequently stimulated with BMPs and/or flow. After 45 minutes, we fixed the cultures and fluorescently labeled the EC nuclei and pSMAD1/5/9 (Fig. 1A).

To quantify responses to ALK1 ligands and SS at single-cell resolution, we developed an image analysis pipeline to measure relative pSMAD1/5/9 localization in the nucleus and cytoplasm as a proxy for BMP/ALK signaling. We collected confocal stacks in two fluorescence channels (DAPI for nuclei; rhodamine for pSMAD1/5/9) and projected the stacks into a single maximum intensity projection image. Segmented ROIs of nuclei from the DAPI channel allowed us to quantify levels of pSMAD1/5/9 inside and outside the nucleus and calculate an “ALK signaling index” (ASI). We defined the ASI as the fraction of total pSMAD1/5/9 found within nuclei (Fig. 1B & Supplementary Fig. 1; see Methods for details). This pipeline, combined with our multimodal microfluidic device, allows ECs to be subjected to a range of ligands and flow conditions and observed with a common imaging framework. The ASI measured with this approach allows unbiased quantitation of EC responses at the single-cell level. We use this device and the automated image analysis pipeline throughout this paper.

### SS increases the sensitivity of ALK1 to BMP9, BMP10 and BMP9/10

To understand how ALK1 ligands and SS interact to stimulate EC signaling, we quantified the ASI in human umbilical vein ECs (HUVECs). First, we tested whether SS interacts with added serum (Fig. 2A), which contains BMP9, BMP10, and BMP9/10 [54]. Without serum or SS, the ASI represents the lowest detectable levels of ALK signaling as it approaches the noise floor of our image acquisition pipeline. The noise floor of the ASI manifests as similar levels of pSMAD1/5/9 fluorescence in the cytoplasm and nucleus, yielding an ASI of ∼ 0.5 (Fig. 1B). HUVECs treated with 2% serum or exposed to SS (40 dyn/cm^2^) increased ASI to between 0.55 and 0.6. When cells were exposed to both 2% serum and SS, ASI increased to about 0.75. The increase in ASI of HUVECs exposed to both serum and SS demonstrates that ALK signaling increases synergistically, beyond the simple addition of responses to each stimulus alone.

**Figure 2.**
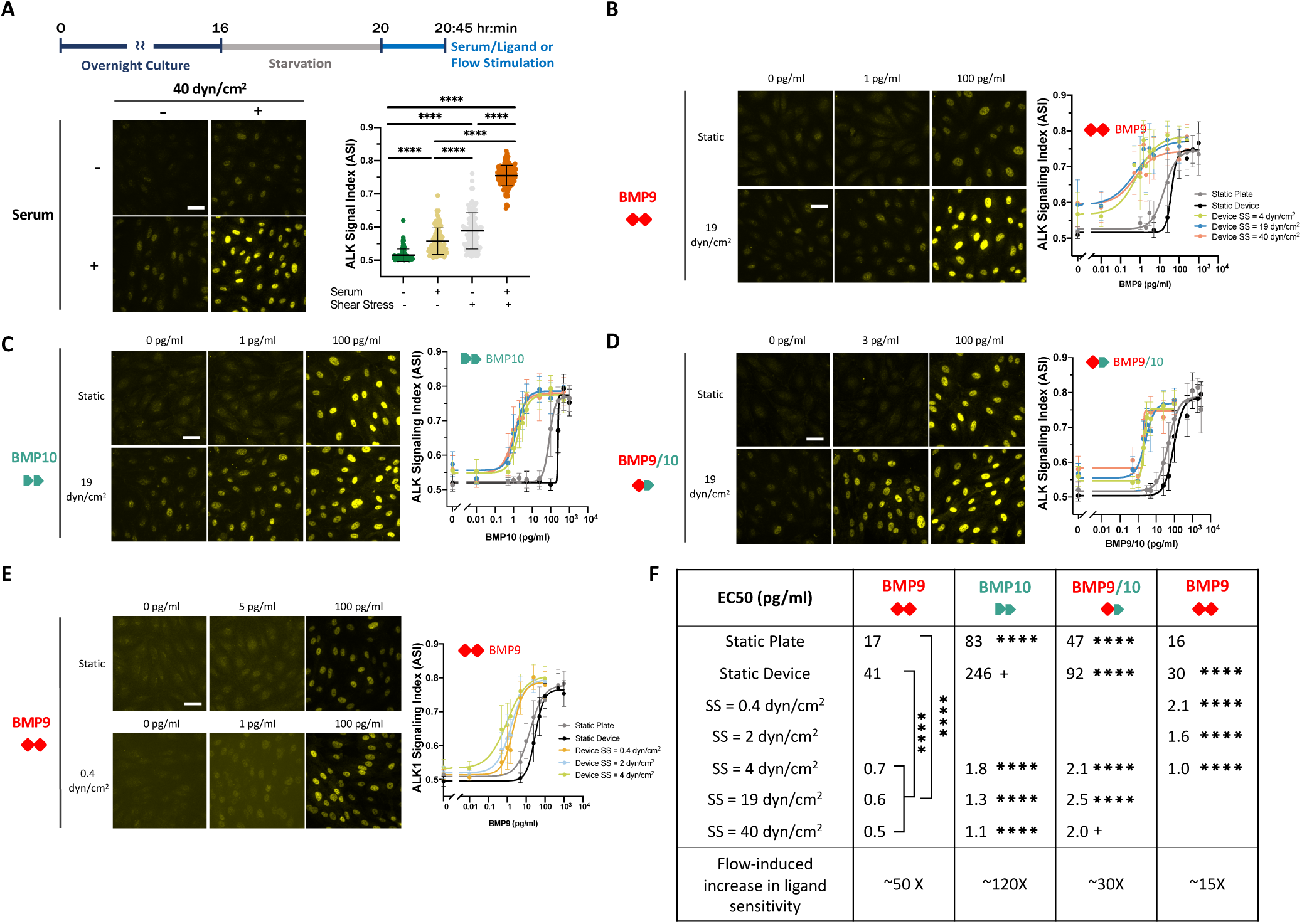
Shear stress synergizes with ALK1 ligands to induce nuclear pSMAD1/5/9 in ECs. (A) Confluent HUVECs were serum starved in 0.2% BSA medium for 4 hours, refreshed with 0.2% BSA medium or 2% serum in EBM2, and cultured under static (in culture plate) or flow (40 dyn/cm^2^ SS) conditions for 45 minutes. Fixed cells were stained for pSMAD1/5/9 and ALK signaling index (ASI) was calculated as per methods. (B-D) Confluent HUVECs were serum starved in 0.2% BSA medium for 4 hours then exposed for 45 minutes to increasing concentrations of BMP9 (B), BMP10 (C) or BMP9/10 (D) in a static culture plate (gray), or in the microfluidic device under static (black) or flow conditions: 4 dyn/cm^2^ (green), 19 dyn/cm^2^ (blue) or 40 dyn/cm^2^ (orange). (E) Confluent HUVECs were serum starved in 0.2% BSA medium for 4 hours then exposed for 45 minutes to increasing concentrations of BMP9 in static culture plate (gray), or in the microfluidic device under static (black) or flow conditions: 0.4 dyn/cm^2^ (orange), 2 dyn/cm^2^ (blue) or 4 dyn/cm^2^ (green). (F) Summary table, ligand EC50s (pg/mL). EC50s under static (plate, device) and flow conditions were averaged to calculate the flow-induced increase in ligand sensitivity. Data were collected from three independent experiments. Two-way ANOVA with Tukey test for multiple comparisons. EC50s were compared across lines in the same column using the extra sum-of-squares F test; **** p < 0.0001, + unable to calculate the comparison of EC50. Scale bar = 40 µm.

We next sought to assay the interaction between specific ALK1 ligands and SS (Fig. 2B-F). We seeded HUVECs in our microfluidic device and treated them with increasing concentrations of three ALK1-specific ligand dimers—BMP9, BMP10 and the heterodimer of BMP9 and 10 (BMP9/10)—under static or flow conditions and measured the ASI in all three uniform SS regions, corresponding to 4, 19, and 40 dyn/cm^2^. We generated 8-point dose-response curves and calculated ligand concentrations that generate a half-maximal response (EC_50_). As previously reported [26], SS dramatically enhanced sensitivity to BMP9, reducing the BMP9 EC_50_ from ∼17 pg/mL (culture plate) or 41 pg/mL (device) under static conditions to 0.5-0.7 pg/mL with flow (Fig. 2B,F). In other words, flow induced an approximately 50-fold increase in HUVEC sensitivity to BMP9 (based on averaged EC_50_ for static and flow conditions). Similarly, we show for the first time that SS enhanced HUVEC sensitivity to BMP10 and BMP9/10 by 120- and 30-fold, respectively (Fig. 2C-D,F). The downward shift in BMP9 EC_50_ was independent of the SS magnitude, whereas the decrease in BMP10 EC_50_ was modestly correlated with an increase in SS magnitude (Fig. 2B-D,F).

### Flow-ALK1 ligand synergy is detected at very low SS magnitudes

In our initial experiments (Fig. 2A-D), HUVECs were exposed to SS from 4 to 40 dyn/cm^2^, a range that includes physiological flows encountered in most mature veins and arteries [55]. However, developing embryonic blood vessels experience lower SS magnitudes: for example, ∼0.4 - 5.5 dyn/cm^2^ in embryonic mice [56]. Therefore, we next investigated responses of HUVECs to BMP9 at low SS magnitudes of 0.4, 2, and 4 dyn/cm^2^. The EC_50_ of BMP9 shifted from ∼16-30 pg/mL under static conditions to 1-2 pg/mL with flow (Fig. 2E,F), with a SS magnitude-dependent increase in sensitivity between 0.4 and 4 dyn/cm^2^. These data demonstrate that flow-ALK1 ligand synergy occurs at extremely low magnitudes of SS that ECs might encounter in the developing vasculature.

### SS induces ALK1 signaling in the presence of trace levels of ALK1 ligand

Our HUVEC ALK1-ligand dose response curves revealed that the ASI under SS was moderately higher than the static condition, even in the absence of added serum or ligand (Fig. 2). To further investigate this effect, we measured the ASI in HUVECs incubated in 0.2% BSA medium and exposed to increasing SS ranging from 0.05 to 40 dyn/cm^2^ (Fig. 3). Interestingly, the ASI exhibited a sigmoidal response to SS (Fig. 3A) that saturated at 19 dyn/cm^2^. The minimum SS magnitude that increased the ASI was 0.4 dyn/cm^2^ with a half-maximal response to flow (EF_50_) at 2.8 dyn/cm^2^.

**Figure 3.**
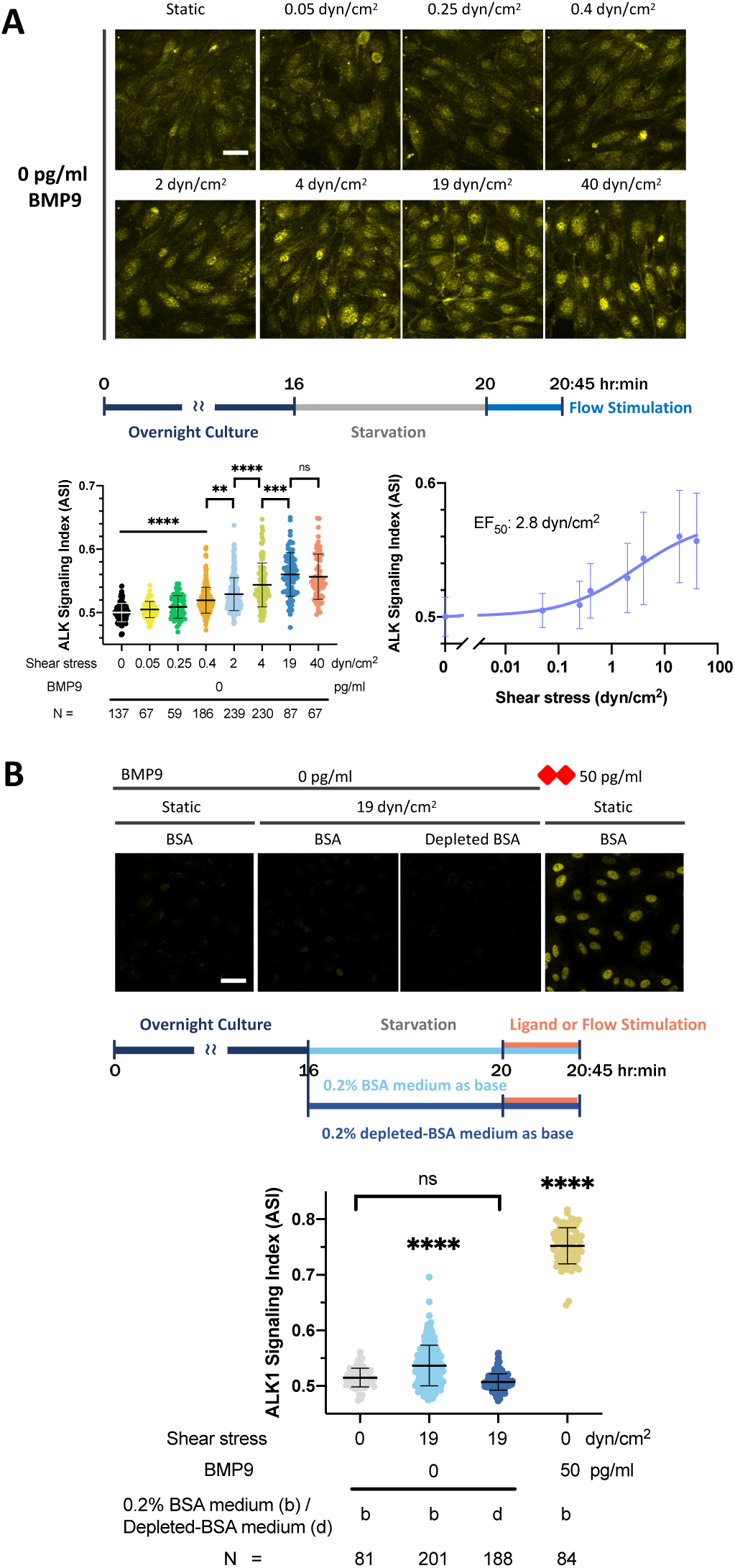
ALK1 signaling in ECs is activated by very low SS magnitude in an ALK1 ligand-dependent manner. (A) HUVECs were serum starved in 0.2% BSA medium for 4 hours then exposed for 45 minutes to static conditions (in culture plate) or SS at the magnitudes indicated. Fixed cells were stained for pSMAD1/5/9 and ASI was calculated as per methods. **(B)** HUVECs were serum starved in 0.2% BSA medium (b) or 0.2% ALK1-ligand depleted-BSA medium (d) for 4 hours followed by a medium refresh with the same medium. Cells were then exposed for 45 minutes to static conditions (in culture plate) or 19 dyn/cm^2^ SS (in device). Stimulation with 50 pg/mL BMP9 was performed as a positive control. Fixed cells were stained for pSMAD1/5/9 and ASI was calculated as per methods. Data in (**A**) were collected from 3 independent experiments; data in (**B**) show one representative trial of 3 independent experiments. N = number of ECs analyzed per group. One-way ANOVA and Tukey multiple comparison test. ** p = 0.009, *** p = 0.0003, **** p < 0.0001. Note: ASI axis scales differ in (**A**) and (**B**). Scale bar = 40 µm.

This low level of signaling, detectable with our sensitive image analysis pipeline, suggested that flow alone stimulates nuclear accumulation of pSMAD1/5/9 equivalent to femtogram/mL levels of ALK1 ligand (Fig. 2B-D). To determine whether this signal response without added ligand was due to trace levels of ALK1 ligand in our commercial source of BSA, we pre-treated the BSA with ALK1-Fc, a soluble chimeric protein consisting of the extracellular BMP-binding domain of ALK1 fused to the Fc portion of IgG, to deplete any residual ALK1 ligands [18]. We removed the ALK1-Fc complex and used 0.2% ALK1-Fc-depleted-BSA in basal media. ALK1-Fc-depletion of BSA medium eliminated the flow-alone response, with ASI decreasing from 0.53 to 0.50. From this shift, we estimate that our 0.2% BSA medium contains ultra-low levels (femtogram/mL) of ALK1 ligand (Fig. 3B). For the remainder of the paper, we will refer to the signal response in the presence of flow but without added ligand as an ultra-low ligand response.

### The ultra-low ligand response and ligand/flow synergy effect are ALK1 receptor-dependent

We next wanted to determine whether the ultra-low ligand response (0 pg/mL BMP9) and the ligand/flow synergy effect (1 pg/mL BMP9) were dependent on ALK1 (Fig. 4A). Accordingly, we knocked down *ALK1* in HUVECs (ALK1-KD) using lentiviral-transduced *ALK1* shRNA, incubated KD and control cells either without added ligand or with BMP9, and analyzed ASI under static or 19 dyn/cm^2^ SS conditions. We achieved 90 to 95% *ALK1* mRNA knockdown and ∼ 85% ALK1 protein knockdown by 6 days post-transduction (Supplementary Fig. 2). With ALK1-KD, the flow response was significantly blunted without (left) and with added ligand (right), demonstrating that both the ultra-low ligand response and ligand/flow synergy effect depend on the ALK1 receptor (Fig. 4B).

**Figure 4.**
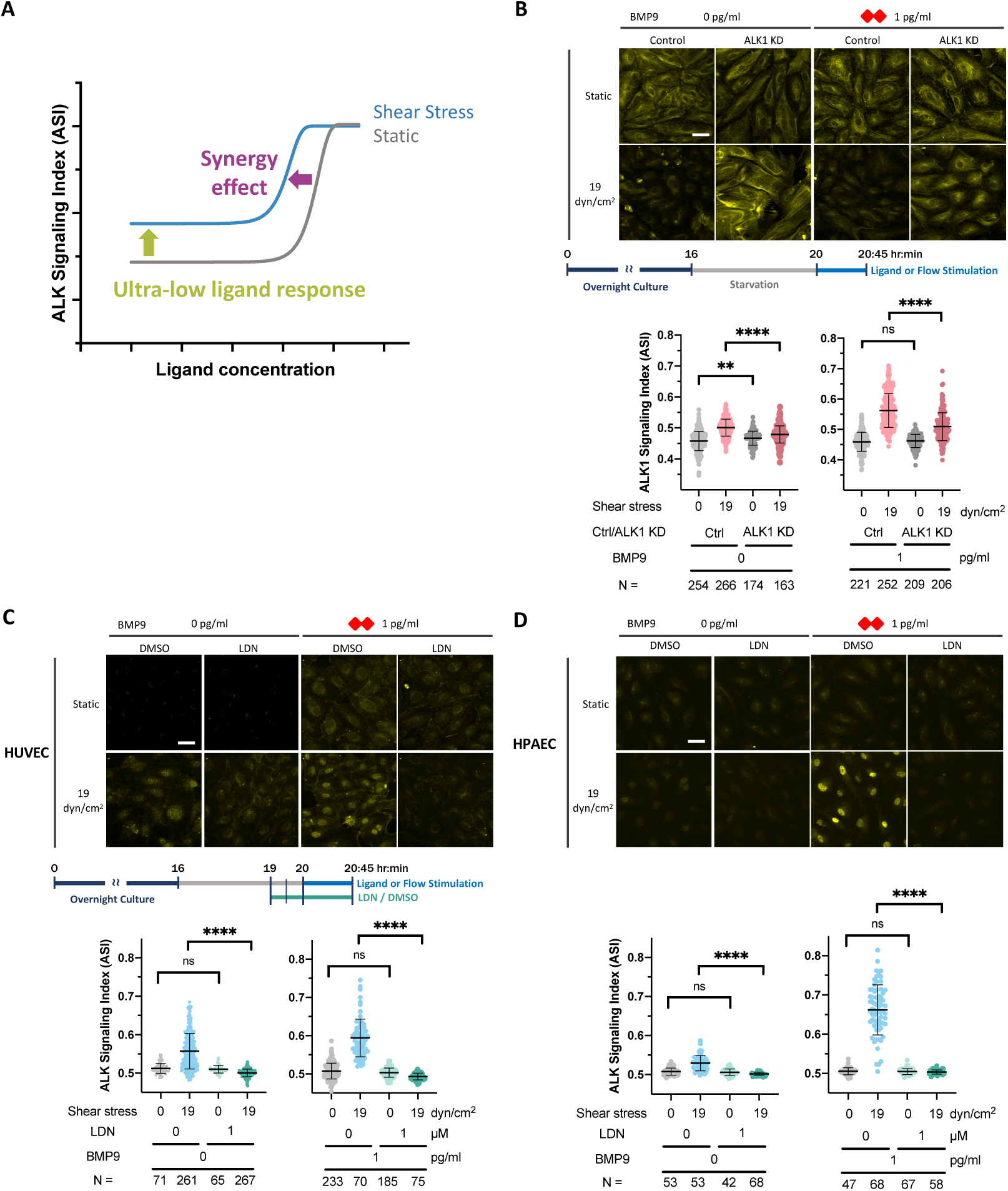
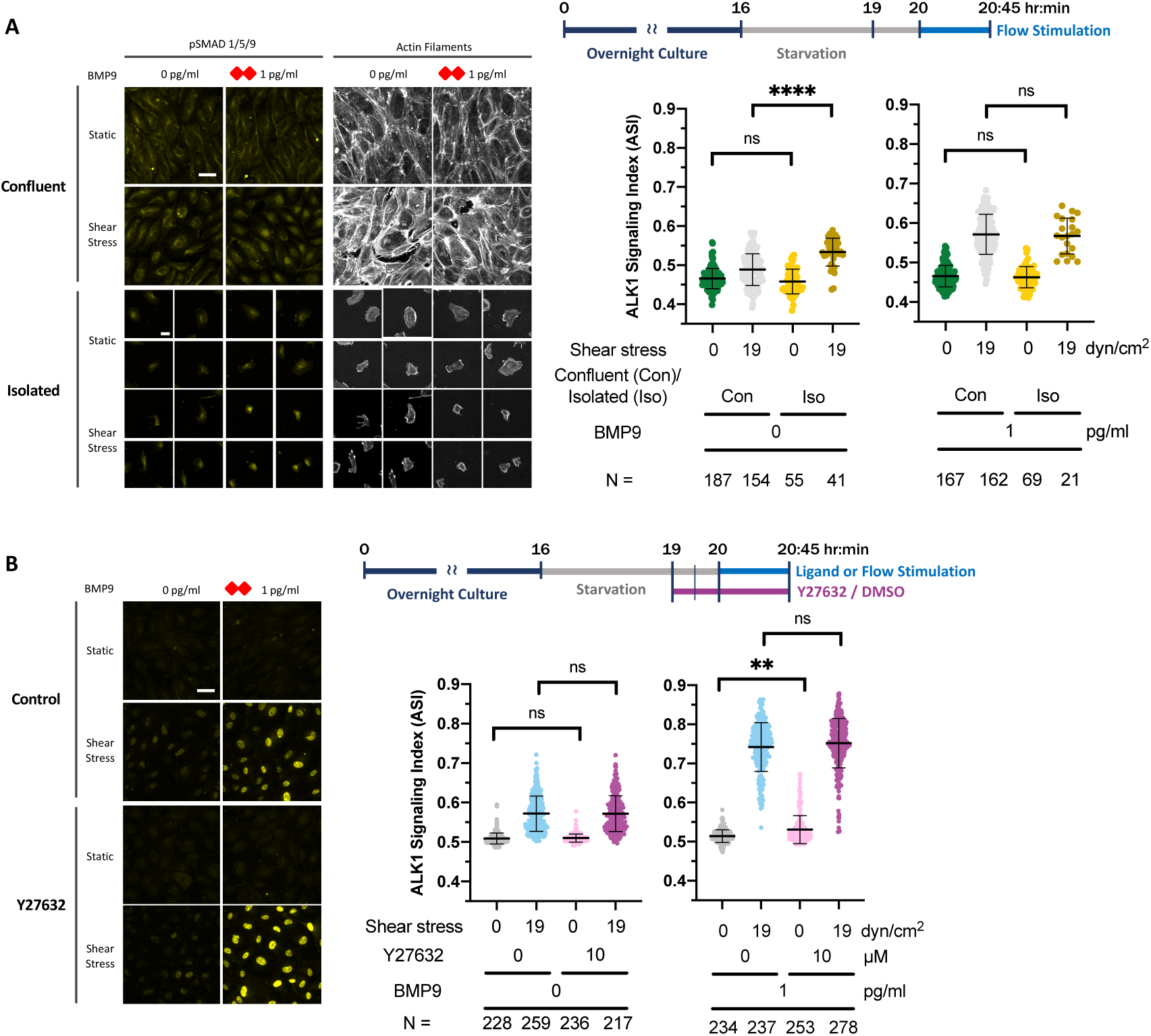
ALK1 protein expression and ALK activity are required for the ultra-low ligand response and synergy effect. (**A**) Schematic depicting the ultra-low ligand response (effect of flow without added ligand) and the synergy effect (effect of flow with added ligand). (**B**) Control and ALK1 KD HUVECs were serum starved in 0.2% BSA medium for 4 hours followed by a medium change to fresh 0.2% BSA medium, without (left) or with (right) 1 pg/mL BMP9, then exposed for 45 minutes to static conditions (culture plate, 0 dyn/cm^2^) or SS (19 dyn/cm^2^). Fixed cells were stained for pSMAD1/5/9 and ASI was calculated as per methods. (**C-D**) HUVECs (**C**) or HPAECs (**D**) were serum starved in 0.2% BSA medium for 4 hours. LDN or DMSO was added 3 hours into the starvation period, followed by treatment without (left) or with (right) 1 pg/mL BMP9. Cells were exposed for 45 minutes to static conditions (culture plate, 0 dyn/cm^2^) or SS (19 dyn/cm^2^), fixed and stained for pSMAD1/5/9, and ASI calculated as per methods. (**B,C**) Data were combined from three independent experiments. (**D**) Data are representative of one trial of three independent experiments. N = number of ECs analyzed per group. One-way ANOVA and Tukey multiple comparison test. ** p = 0.0054, **** p < 0.0001. Scale bar = 40 µm.

To determine whether these responses required ALK1 kinase activity, we used a pharmacological type I receptor kinase inhibitor (LDN193189, or LDN) known to inhibit ALK1, ALK2 (ActRIA), ALK3 (BMPRIA) and ALK6 (BMPR1B) [57, 58]. We administered LDN to HUVECs 60 min before flow stimulation and during the following 45 min of flow at 19 dyn/cm^2^. Both the ultra-low ligand response (left) and the synergy effect (right) were eliminated by LDN (Fig. 4C). Given the ASI dependence on *ALK1* expression, this result supports the role of ALK1 kinase activity in SS sensing.

To test whether the ultra-low ligand response and synergy effects and their dependence on ALK1 were specific to venous ECs, we repeated the LDN experiment with primary human pulmonary artery endothelial cells (HPAEC). Similarly to HUVECs, HPAECs exhibited both ultra-low ligand response (left) and the ligand/synergy effect (right), which were both eliminated by LDN (Fig. 4D). These data indicate that both venous and arterial ECs can mount a synergistic response to a combination of ALK1 ligands and flow.

### Requirements for mechanosensing in the flow-ALK1 ligand synergy effect

#### 1. Mechanosensing via cell-cell adhesion is not required for flow-ALK1 ligand synergy in HUVEC

Our data thus far demonstrate that SS sensitizes HUVECs to respond to trace levels of BMP ligands in an ALK1-dependent manner. To elucidate the molecular mechanisms responsible for flow mechanosensing in both ultra-low ligand response and synergy effect, we first tested the requirement for cell-cell junctions. It has been proposed that cell-cell junctions sense flow and are required for subsequent cell alignment. Elements important in this mechanotransduction complex include VE-cadherin, VEGFR2, and PECAM-1 [11], and tension at VE-cadherin in HUVEC cell-cell junctions changes following the onset of flow [59]. Therefore, we hypothesized that ultra-low ligand response and flow-ligand synergy effect require endothelial sheet integrity and the formation of tension-transmitting junctions. To test this hypothesis, we compared the ASI between confluent and isolated HUVECs subjected to 19 dyn/cm^2^ SS, with and without BMP9. Surprisingly, both isolated and confluent ECs behaved similarly in these assays (Fig. 5A, and Supplementary Table 1). These results indicate that both responses to SS are cell-cell junction independent.

**Figure 5.**
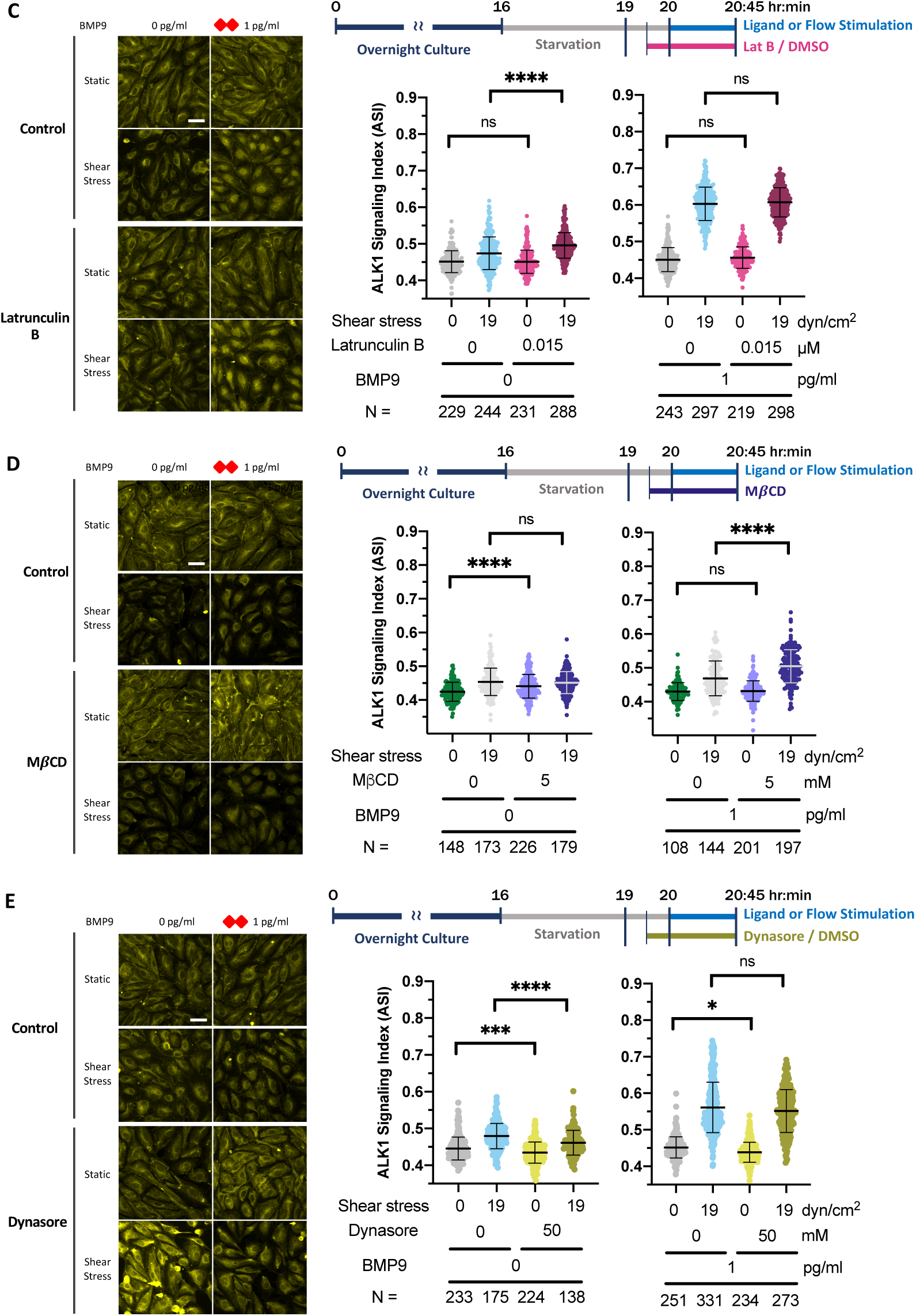
Requirements for EC junctions, actin cytoskeleton, cholesterol-rich microdomains, and endocytosis in the ultra-low ligand response and flow/ligand synergy effect. (A) Confluent or isolated HUVECs were serum starved in 0.2% BSA medium for 4 hours then exposed for 45 minutes to static (culture plate, 0 dyn/cm^2^) or SS (19 dyn/cm^2^) conditions, without or with 1 pg/mL BMP9. Fixed cells were stained for pSMAD1/5/9 and ASI was calculated as per methods. (B-E) Confluent HUVECs were serum starved in 0.2% BSA medium for 4 hours with Y273632 (B), latrunculin B (C), MβCD (D) or dynasore (E) added at indicated times/concentrations during the starvation period, then exposed for 45 minutes to static conditions (culture plate, 0 dyn/cm^2^) or SS (19 dyn/cm^2^), with or without 1 pg/mL BMP9. Fixed cells were stained for pSMAD1/5/9 and ASI was calculated as per methods. N = number of ECs analyzed per condition. Data within the same BMP9 concentration were analyzed by two-way ANOVA and Tukey test for multiple comparisons; ns = no significant difference, *p = 0.043, **** p < 0.0001. Scale bar = 40 µm.

#### 2. Mechanosensing via the cytoskeleton does not contribute to flow-ALK1 ligand synergy in HUVEC

Early studies of EC responses to SS proposed that mechanosensing occurs via tension acting on the cytoskeleton, which subsequently alters cell behaviors that drive cell alignment [1]. To test the role of cytoskeletal-dependent mechanosensing in the ultra-low ligand response and flow-ligand synergy effect, we inhibited actomyosin contractility using the p160 Rho kinase (ROCK) inhibitor, Y27632 [59]. We validated drug efficacy in HUVECs by demonstrating that treatment with 10 mM Y27632 for 60 minutes enhanced junctional tortuosity, an indication of decreased contractility [49] (Supplementary Fig. 3). We next compared the ASI between vehicle and Y27632-treated HUVECs subjected to 19 dyn/cm^2^ SS, with or without added BMP9. However, neither the ultra-low ligand response nor the ligand/flow synergy effect was significantly decreased when cytoskeletal contractility was disrupted (Fig. 5B and Supplementary Table 1).

As an orthogonal method to test the importance of the cytoskeleton, we reduced F-actin levels by inhibiting actin polymerization with latrunculin B [50]. To maintain normal EC morphology under 19 dyn/cm^2^, HUVECs were treated with latrunculin B at a relatively low dose (0.015 µM), resulting in a reduction of apical actin while leaving basal actin unaffected (Supplementary Fig. 4). We compared the ASI between vehicle and latrunculin B-treated HUVECs subjected to 19 dyn/cm^2^ SS, with or without added BMP9. As with Y27632 treatment, neither the ultra-low ligand response nor the ligand/flow synergy effect was significantly decreased by latrunculin (Fig. 5C and Supplementary Fig. 5). Thus, we conclude that flow-enhanced ALK1 signaling does not require cytoskeletal-mediated mechanosensing pathways.

#### 3. Mechanosensing via cholesterol-enriched microdomains does not contribute to flow-ALK1 ligand synergy in HUVEC

Cholesterol-enriched microdomains in the plasma membrane reflect a preferential association between specific lipids and proteins and have been implicated in cell surface receptor-mediated signal transduction [60]. Accordingly, we hypothesized that flow may alter cholesterol-enriched microdomains leading to ALK1/type II receptor clustering and, consequently, enhanced ligand-mediated signaling and increased pSMAD1/5/9 nuclear localization. To test this hypothesis, we disrupted cholesterol-enriched microdomains with methyl-β-cyclodextrin (MβCD), which chelates cholesterol from cellular lipid bilayers and decreases signaling transduction via cholesterol-enriched microdomains [61–63].

To validate this drug efficacy, we treated HUVECs with 5 mM MβCD and observed reduced numbers of cholera toxin B-subunit (CTxB) puncta, which is endocytosed and transported to the endoplasmic reticulum [64], indicating effective disruption of cholesterol-enriched microdomains (Supplementary Fig. 6). We next compared the ASI between vehicle and 5 mM MβCD-treated HUVECs subjected to 19 dyn/cm^2^ SS, with or without added BMP9. Neither the ultra-low ligand response nor the ligand/flow synergy effect was significantly decreased (Fig. 5D and Supplementary Fig. 5). These data suggest that flow-enhanced ALK1 signaling does not require cholesterol-enriched microdomains.

#### 4. Endocytosis contributes to the ultra-low ligand response

Receptor clustering can activate signaling pathways involving cell surface receptors [65]. Although most receptor clustering is triggered by ligand binding, endocytosis can aid in colocalizing or clustering receptors [66–69]. Thus, we hypothesized that flow may enhance endocytosis, thereby clustering ALK1 receptors and increasing pSMAD1/5/9 signaling. Considering that dynamin-2 is crucial for ALK1 endocytosis [70], we used dynasore, a potent and specific dynamin1/2 inhibitor, to interfere with endocytosis [71]. We compared the ASI between vehicle and 100 µM dynasore-treated HUVECs subjected to static conditions or 19 dyn/cm^2^ SS, with or without added BMP9. HUVECs treated with dynasore exhibited a reduction in ASI under static and flow conditions in the absence of added ligand, suggesting that the ultra-low ligand effect requires endocytosis. By contrast, although dynasore modestly decreased the ASI in response to 1 pg/mL BMP9 under static conditions, it had no effect on the flow-BMP9 synergy effect (Fig. 5E and Supplementary Fig. 5).

## Discussion

### Flow/ligand synergy is common to all ALK1 ligands and observed at very low SS magnitudes and ligand concentrations

Our microfluidic system allows precise stimulation of EC cultures via specific ALK1 ligands and SS cues. Single-cell analysis allows us to account for cell-to-cell variation and flow conditions along the 31-mm length of the channel through three different laminar SS regions. From these analyses, we confirmed the synergy effect between SS and the ALK1 ligand BMP9 [26](Fig. 6A), and extended this finding to two additional ALK1 ligands, BMP10 and BM9/10. Additionally, we identified two previously undescribed interactions between ALK1 ligand and flow sensing. The first is an ultra-low ligand response stimulated by SS in the presence of trace (femtogram/mL) levels of ALK1 ligand (Fig. 3A). This ultra-low ligand response is ALK1-dependent and SS-magnitude dependent and displays a sigmoidal dose-response between 0.4 and 19 dyn/cm^2^ with an EF_50_ of 2.8 dyn/cm^2^. The second novel response is a strong synergy between ligand and even very low SS magnitudes (Fig. 4A). With added ligand, SS magnitudes ≥ 0.4 dyn/cm^2^ are sufficient to enhance ALK1 receptor sensitivity, lowering the BMP9 EC_50_ 15-30-fold, with only modest changes with increasing SS magnitude. With the sensitivity of our ASI quantification, we found that the SS threshold for synergy is much lower than previously reported for other examples of flow responses in ECs [26].

**Figure 6.**
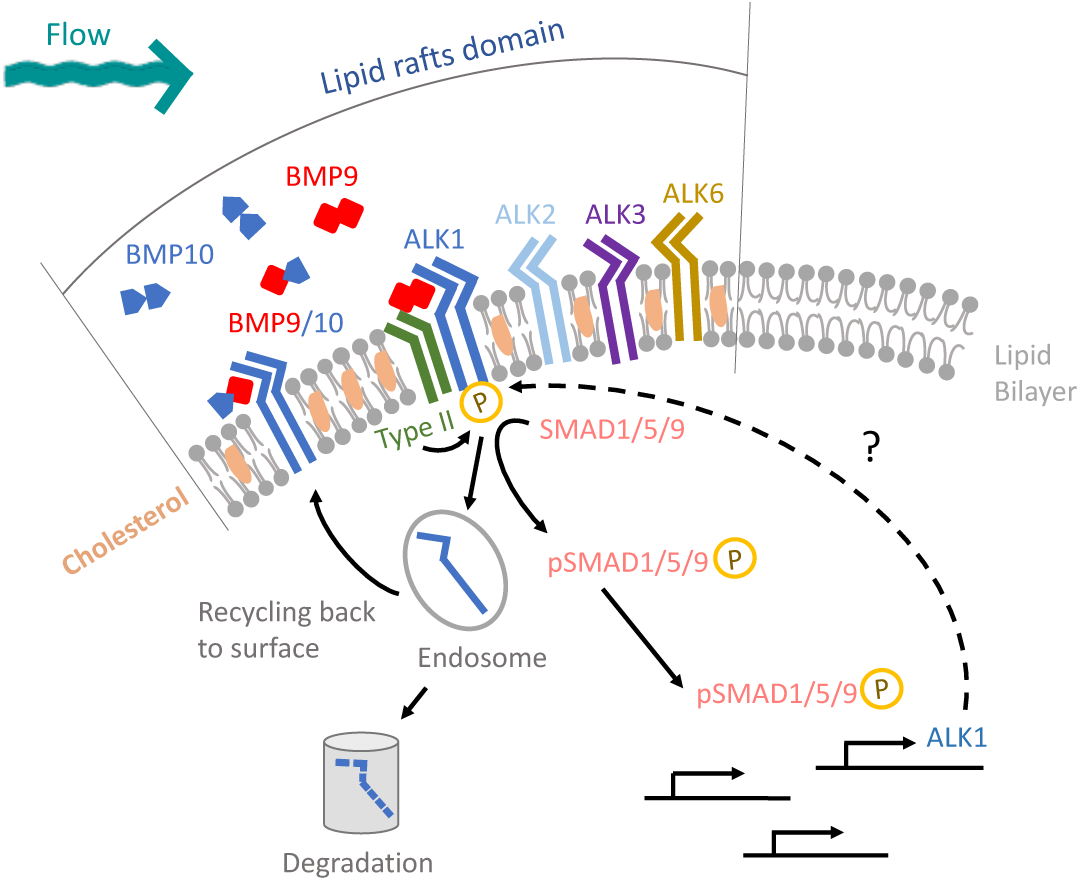
Working model of how fluidic mechanical and chemical cues integrate to enhance ALK1 signaling.

### Testing potential mechanisms of mechanosensing through sensitive analysis of pSMAD1/5/9 nuclear localization

To understand how fluid-mechanical cues are sensed and then synergize ALK1 signaling, we developed a working model that integrates multiple mechanosensing pathways. Formally, each of these pathways may be operating simultaneously and, to our knowledge, none are mutually exclusive. To test mechanosensing mechanisms that might lead to enhanced pSMAD1/5/9 signaling (Fig. 6), we devised a series of tests. (1) First, we hypothesized that endothelial sheet integrity and cell-cell junctions are required for both types of flow sensing. By comparing the ASI between confluent and isolated cells, we found that ultra-low ligand response and flow-synergy effect were unchanged in isolated cells (Fig. 4A), suggesting that both effects are cell-cell junction independent. (2) Second, we tested whether the cytoskeleton-mediated mechanosensing pathways were responsible for flow/ALK1 sensing. We inhibited myosin contractility and F-actin polymerization by using Y27632 and latrunculin B, respectively. However, neither treatment altered the flow-ligand interactions, ruling out a cytoskeletal role for mechanosensing required for synergy. (3) Third, we investigated the role of cholesterol-enriched microdomains. Our results suggest that cholesterol-enriched microdomains and caveolae-dependent endocytosis do not play a role in either the SS-mediated synergy or the ultra-low ligand response. (4) Finally, we explored the role of dynamin-dependent endocytosis. We used the dynamin inhibitor dynasore and found that ECs maintained flow/added ligand synergy but demonstrated a reduced response to ultra-low ligand. This suggests that dynamin-dependent endocytosis contributes to the mechanosensing pathway responsible for the ultra-low ligand response but not the synergy effect.

### ALK1-mediated mechanosensing is regulated via various pathways of endocytosis in a context dependent manner

Intracellular membrane trafficking is likely to play an important role in modulating ALK1 signaling and synergy under SS [72]. EC barrier functions are sensitive to large disruptions in membrane trafficking, such as intercellular junctional permeability [73]. Therefore, we sought pharmacological inhibitors with short-term and acute impact. Since dynamin-2 inhibits caveolae internalization in a GTPase-independent manner, we assume dynasore would not affect caveolae internalization [74]. Therefore, our work using these inhibitors suggests that ALK1 internalization through clathrin-dependent endocytosis significantly decreases in both static and flow conditions with no added ligand. Our results agree with a recent study demonstrating that ALK1 undergoes rapid clathrin-dependent endocytosis independent of added ligand [75]. By contrast, other studies have suggested that BMP9-mediated ALK1 signaling involved caveolae/caveolin-1-dependent endocytosis [44, 76, 77]. These discrepancies may be due to differences in both BMP9 and SS stimulation intensity and duration. Prior studies used saturating doses of BMP9 (100 pg/mL to 10 ng/mL) under static conditions [70], while others exposed cells to low SS over longer times (1 dyn/cm^2^ for 24 hours) [44]. Additionally, these studies offered a limited readout of ALK1 signaling, utilizing pooled analysis of cellular pSMAD1/5/9 signaling level via western blot and RNA-sequencing dataset. Our work tested synergy between flow and BMP9 at much lower doses (0 pg/mL and 1 pg/mL) with a 45-minute stimulation period and involved a more sensitive readout of AK1 signaling, utilizing immunofluorescence with single cell resolution. As a result, we detected enhanced ALK1 signaling requiring clathrin-dependent endocytosis after 45 minutes of flow. Combined with previous work, our results suggest context-dependent mechanisms of ALK1 internalization upon stimulation; we speculate that the basal and low dose BMP9/ALK1 activation signals via clathrin-dependent endocytosis, while long-term exposure to saturating doses may signal via caveolae/caveolin-1 dependent endocytosis. Further work using genetic or optogenetic interrogation of endocytic pathways with live reporters of ALK1 signaling will be required to confirm and test pathways leading to ALK1 internalization and signaling at the onset of flow.

Membrane transport can play a broad role in modulating ligand-receptor signaling in ECs. For example, VEGFR2 is internalized through the clathrin-mediated pathway in the absence of ligand and mainly through CDC42-mediated macropinocytosis with the application of VEGF-A [78]. Moreover, ALK5, another type I TGFβ receptor (TβRI), appears to be internalized through both clathrin- and caveolin-1-dependent endocytosis. In this case, endocytosed clathrin-coated vesicles and caveolar vesicles appear to fuse, forming a novel type of caveolin-1 and clathrin double-positive endocytic vesicle [79]. Therefore, we hypothesize that low levels of ALK1 receptors may be endocytosed via clathrin-dependent endocytosis under short-term flow conditions without added ligands, and then transition to being internalized via caveolin-1-dependent pathways as ligands bind. Our methods cannot distinguish these complex interactions since dynasore not only inhibits clathrin-dependent endocytosis, but may also disrupt the organization of cholesterol-enriched microdomains [80]. As in the above discussion of endocytosis, dissecting the role of membrane trafficking will require future studies using more precise pathway manipulations and detection of ALK1 signaling from within intracellular membrane subcompartments.

### Perturbations that enhance pSMAD1/5/9 signaling

Our experiments were designed to identify pathways that lowered ALK1 signaling, however, we also found perturbations that enhanced signaling. We observed increases in ASI in both isolated ECs and cells treated with latrunculin B following SS exposure in ultra-low ligand conditions, suggesting that cell-cell adhesion and the F-actin cytoskeleton repress the ultra-low ligand response. ASI was also enhanced in ECs treated with MβCD following SS exposure with 1 pg/mL BMP9, suggesting that cholesterol-enriched microdomain-dependent clustering of receptors represses the flow-ligand synergy effect. The high sensitivity of our analysis method allowed us to detect these unexpected interactions.

### Flow-Synergy of ALK1 signaling regulated by endocytosis may reflect a common mechanism coupling EC responses to flow

Synergy between fluid SS and receptor-meditated signaling is not well understood. Several signaling pathways in ECs are also responsive to SS, including signaling activated by VEGF-A, Notch ligands, chemokine CXCL12 and TGF-β1 [13, 43, 81–85]. However, it is unknown whether SS shifts their signaling response synergistically, or whether these responses are simply additive. For instance, VEGF-A and SS can both drive phosphorylation of VEGFR2 to the same degree [13]. Our findings that low levels of ligand synergize with flow to activate ALK1 suggest a reinterpretation of findings that utilize low-serum media as a baseline for flow studies. These media are likely to contain low levels of ligands that may confound the interpretation of these studies. Furthermore, we suggest carrying out ligand-flow interaction studies at ligand concentrations well below saturating levels, and preferably at the ligand’s EC_50_ under flow, where signal responses are most sensitive to perturbations.

### Conclusions

Sensitive single-cell measurements of ECs exposed to both SS and all known ALK1 ligands reveal that the ALK1 receptor is essential for synergistic effects on SMAD1/5/9 signaling, and that this synergy is apparent at concentrations of ligand and magnitudes of shear stress that are much lower than previously reported. Endocytosis pathways appear to be involved in this mechanosensing pathway but resolving the molecular factors and mechanisms of ALK1 receptor trafficking will require further investigation.

## Acknowledgments

We thank Dr. Douglas Marchuk, Duke University, for the ALK1 antibody. This work was supported by Leonard H. Berenfield Graduate Fellowship (to YC), R01 HL136566 (to BLR), DoD W81XWH-21-1-0352 (to BLR), DoD W81XWH-17-1-0429 (to APH). A. Anzell was supported by T32 HL129964.

## Author contributions

B. L. Roman and L. A. Davidson conceived the initial research idea and jointly guided the project. Y. Cheng conducted and analyzed most of the experiments. A. R. Anzell prepared the shRNA transduced cells and conducted the lipid rafts disruption validation. T. A. Schwartze, C. S. Hinck, and A. P. Hinck prepared and purified the BMP9/10 heterodimer. All authors contributed to the interpretation of the results. Y. Cheng wrote the manuscript with support from all authors. All authors read and approved the final version of the manuscript.

