## Supplementary Material for "Shear stress and very low levels of ligand synergize to activate ALK1 signaling in endothelial cells"

### Supplementary Figures, Tables, and Captions

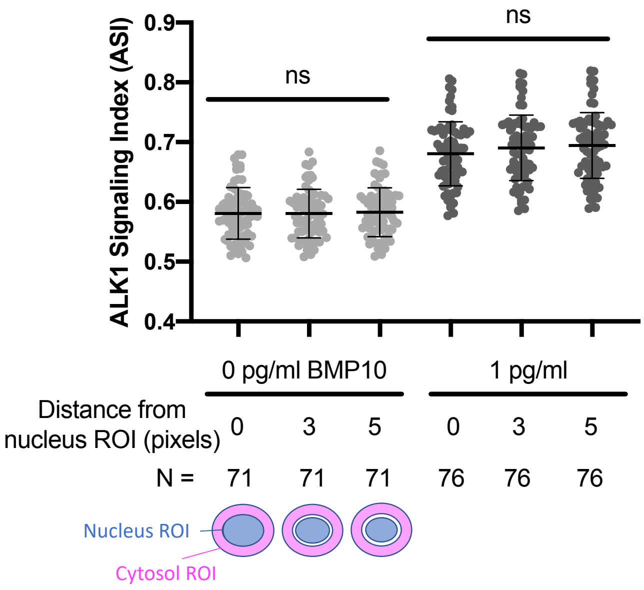


**Supplementary Figure 1 (Supports Figure 1B). Validation of cytosolic region of interest (ROI) used to calculate ALK signaling index (ASI).**
Confluent HUVECs were serum starved for 4 hours in 0.2% BSA medium, treated for 45 minutes with 0 or 1 pg/mL BMP10, then fixed and stained for nucleus (DAPI) and pSMAD1/5/9. To calculate the ASI, the cytosolic ROI was drawn 0, 3, or 5 pixels (0, 1.3, or 2.2 $\mu m$) away from nuclear ROI, and resulting ASIs compared by one-way ANOVA with Tukey multiple comparisons test. ns = not significant. N = 3 independent experiments. Each dot represents 1 EC.

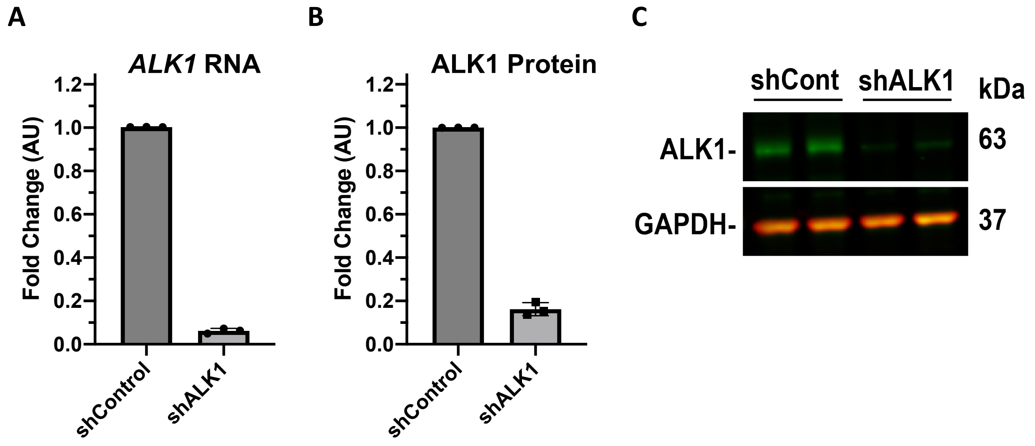


**Supplementary Figure 2 (Supports Figure 4B). Knockdown efficiency of ALK1 in HUVECs.**
HUVECs were transduced with lentivirus carrying control or ALK1-targeted shRNA and analyzed at 6-days post-transduction for *ALK1* mRNA by RT-qPCR (**A**) and ALK1 protein by western blot, normalized to GAPDH (**B**,**C**). N = 3 independent experiments, with duplicate wells (as in **C**) averaged for each N.

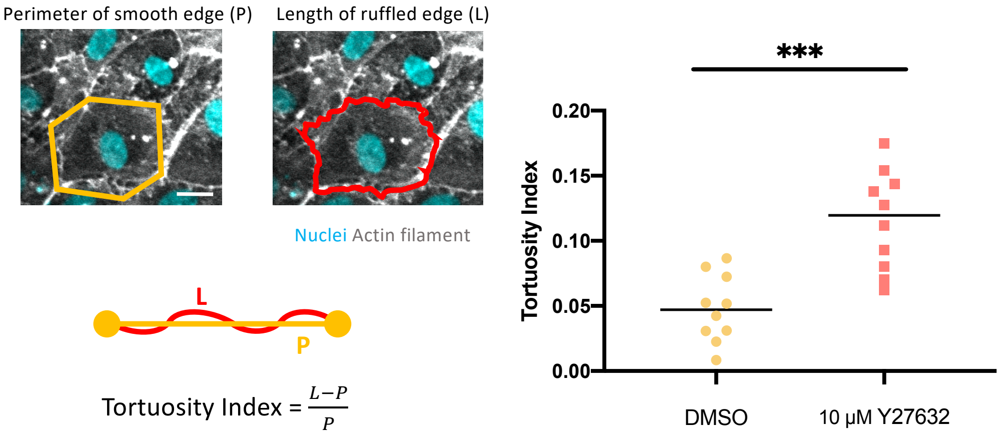


**Supplementary Figure 3 (Supports Figure 5B). Y27632 enhances tortuosity of cell-cell junctions.** Confluent HUVECs treated for 1 hour 45 minutes with 0.1% DMSO or 10 μM Y27632 in 0.1% DMSO**.** Nuclei were labeled with DAPI and the F-actin cytoskeleton was labeled with phalloidin, and the tortuosity index was calculated as indicated. N = 3 independent experiments. Each dot represents 1 EC. Two-tailed unpaired student's t-test. *** p =0.0002 . Scale bar = 20 μm.

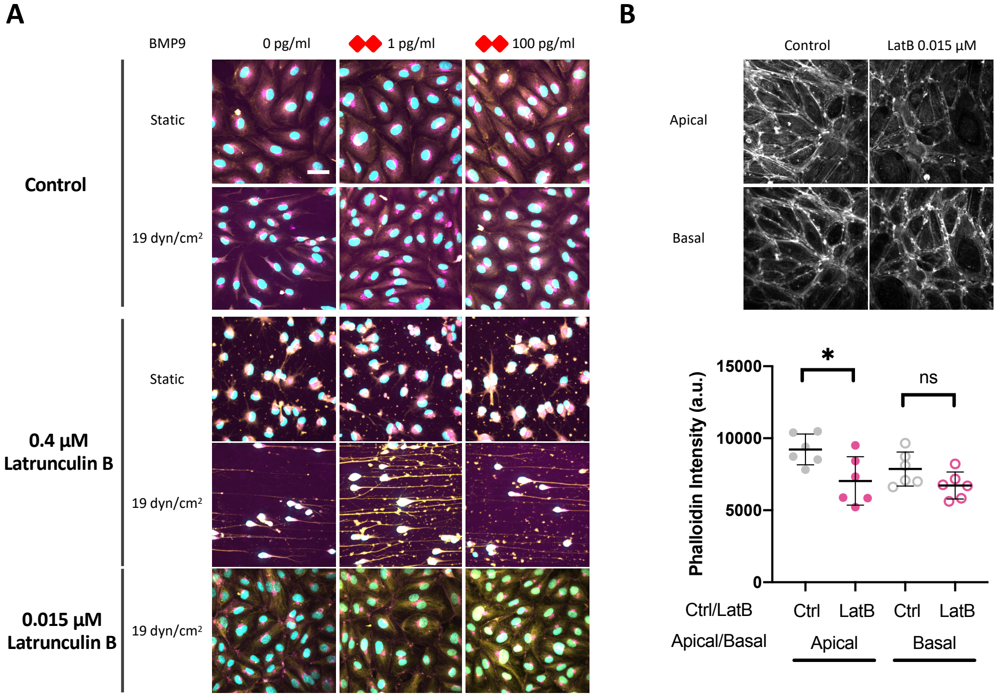


**Supplementary Figure 4 (Supports Figure 5C). HUVECs treated with 0.015 µM Latrunculin B maintain normal EC morphology under shear stress.**
(A) Confluent HUVECs were grown in static conditions or under SS (19 dyn/cm$^2$) and treated with “standard” (0.4 µM) or low (0.015 µM) concentration of latrunculin B, or 0.4% DMSO for 75 minutes. Fixed cells were stained for the nucleus (cyan), pSMAD1/5/9 (yellow), and the Golgi (magenta). (B) Apical actin filament level was decreased with 0.015 µM Latrunculin B treatment, while basal actin remained unchanged under static condition. (stained by Acti-stain 670 phalloidin). Two-tailed unpaired student's t-test, * p = 0.0228. Scale bar = 40 µm.

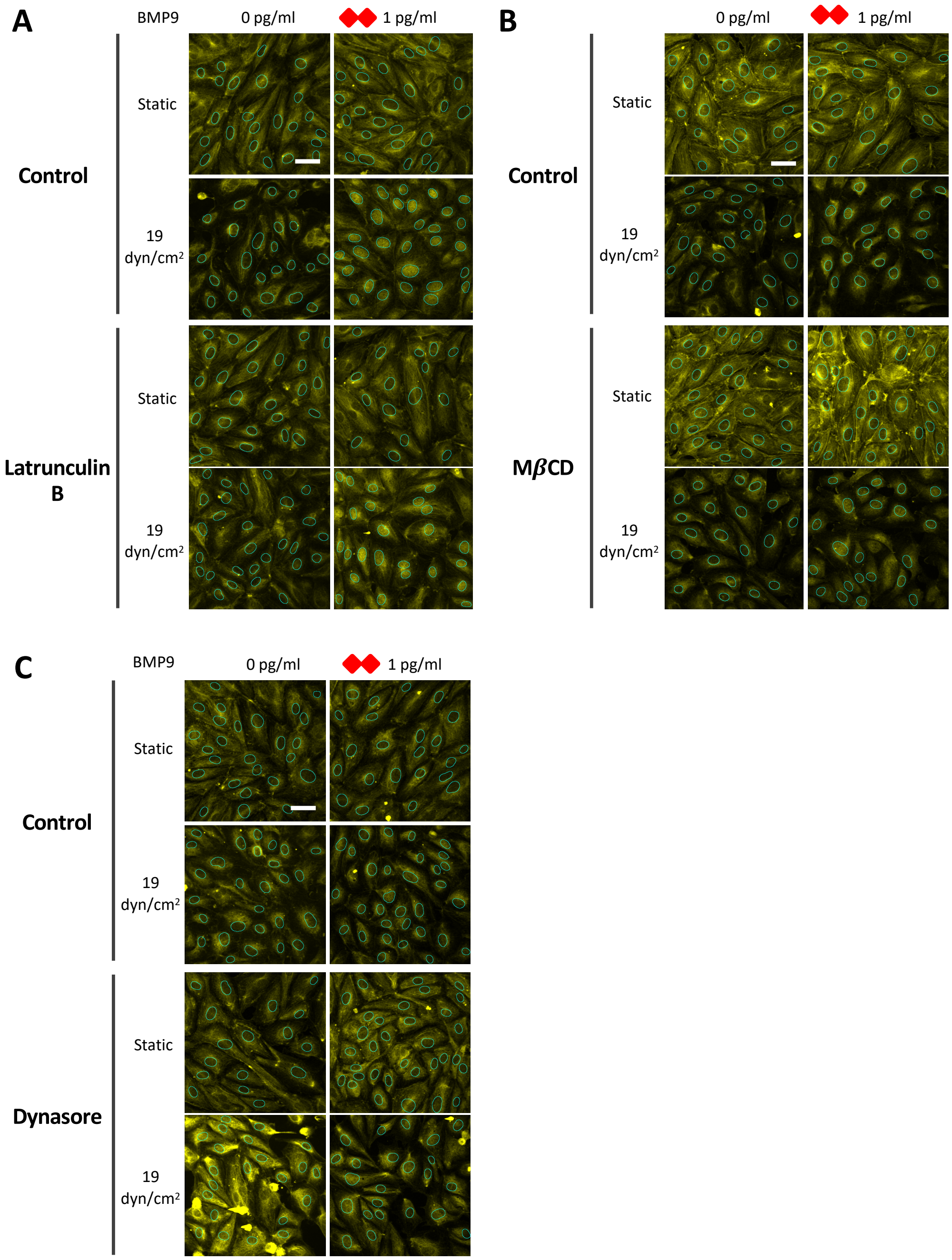


**Supplementary Figure 5 (Supports Figures 5C, 5D, and 5E).**
Nuclear outlines (cyan) and the localization of pSMAD1/5/9 (yellow) in HUVECs treated with latrunculin B (A), MβCD (B) and Dynasore (C). The outlines of nuclei were segmented from the DAPI fluorescence. Scale bar = 40 µm.

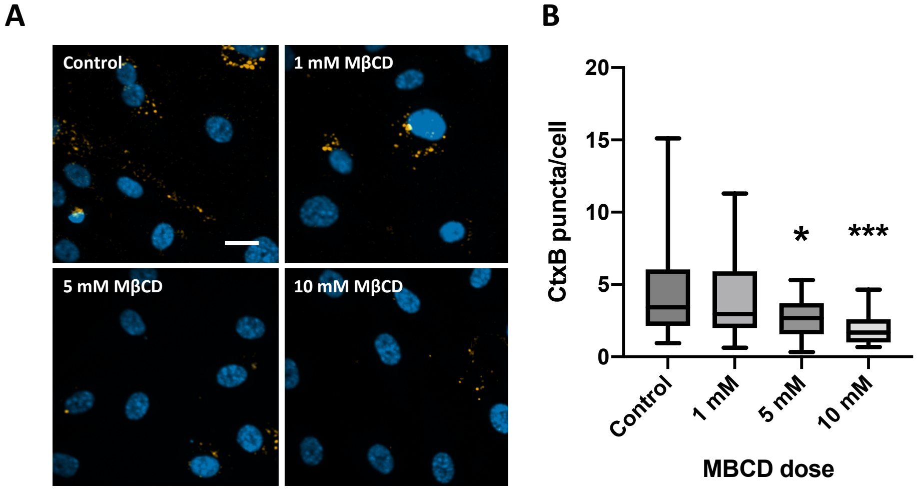


**Supplementary Figure 6 (Supports Figure 5D). MβCD disrupts cholesterol-enriched microdomains.**
HUVECs were treated for 75 minutes with indicated concentrations of MβCD. Fixed cells were stain for nuclei (DAPI) and Cholera toxin subunit B (CtxB), and CtxB puncta counted in 4 fields and N > 250 cells per condition over 3 independent experiments. One-way ANOVA and Tukey post-hoc test, *p = 0.0217; ***p = 0.0005. Scale bar = 20 μm.

**Supplementary Table 1 (Supports Figure 5). Phospho-SMAD1/5/9 mean intensities and standard deviation (SD) under static or flow condition.**

| BMP9 Conc. (pg/ml) | Treatment | Shear Stress (dyn/cm2) | Mean | SD |
|---|---|---|---|---|
| 0 | Confluent | 0 | 0.466 | 0.026 |
| | | 19 | 0.489 | 0.041 |
| | Isolated | 0 | 0.458 | 0.032 |
| | | 19 | 0.534 | 0.036 |
| 1 | Confluent | 0 | 0.465 | 0.027 |
| | | 19 | 0.571 | 0.051 |
| | Isolated | 0 | 0.463 | 0.027 |
| | | 19 | 0.567 | 0.046 |
| 100 | Confluent | 0 | 0.584 | 0.049 |
| | | 19 | 0.648 | 0.040 |
| | Isolated | 0 | 0.552 | 0.031 |
| | | 19 | 0.633 | 0.045 |
| 0 | - | 0 | 0.509 | 0.014 |
| | | 19 | 0.572 | 0.045 |
| | Y27632 | 0 | 0.510 | 0.010 |
| | | 19 | 0.572 | 0.046 |
| 1 | - | 0 | 0.514 | 0.016 |
| | | 19 | 0.742 | 0.062 |
| | Y27632 | 0 | 0.531 | 0.036 |
| | | 19 | 0.752 | 0.063 |
| 100 | - | 0 | 0.761 | 0.040 |
| | | 19 | 0.784 | 0.049 |
| | Y27632 | 0 | 0.779 | 0.034 |
| | | 19 | 0.772 | 0.052 |
| 0 | - | 0 | 0.451 | 0.030 |
| | | 19 | 0.474 | 0.045 |
| | Latrunculin B | 0 | 0.451 | 0.032 |
| | | 19 | 0.496 | 0.035 |
| 1 | - | 0 | 0.451 | 0.033 |
| | | 19 | 0.603 | 0.046 |
| | Latrunculin B | 0 | 0.456 | 0.029 |
| | | 19 | 0.607 | 0.040 |
| 100 | - | 0 | 0.586 | 0.038 |
| | | 19 | 0.647 | 0.041 |
| | Latrunculin B | 0 | 0.609 | 0.038 |
| | | 19 | 0.654 | 0.038 |
| 0 | - | 0.425 | 0.028 | 0.425 |
| | | 0.454 | 0.040 | 0.454 |
| | MβCD | 0.441 | 0.035 | 0.441 |
| | | 0.452 | 0.033 | 0.452 |
| 1 | - | 0.430 | 0.027 | 0.430 |
| | | 0.469 | 0.051 | 0.469 |
| | MβCD | 0.431 | 0.030 | 0.431 |
| | | 0.505 | 0.049 | 0.505 |
| 100 | - | 0.542 | 0.058 | 0.542 |
| | | 0.600 | 0.066 | 0.600 |
| | MβCD | 0.557 | 0.050 | 0.557 |
| | | 0.612 | 0.049 | 0.612 |
| 0 | - | 0 | 0.446 | 0.031 |
| | | 19 | 0.479 | 0.034 |
| | Dynasore | 0 | 0.435 | 0.029 |
| | | 19 | 0.461 | 0.034 |
| 1 | - | 0 | 0.452 | 0.029 |
| | | 19 | 0.561 | 0.069 |
| | Dynasore | 0 | 0.438 | 0.027 |
| | | 19 | 0.552 | 0.059 |
| 100 | - | 0 | 0.540 | 0.044 |
| | | 19 | 0.640 | 0.055 |
| | Dynasore | 0 | 0.556 | 0.039 |
| | | 19 | 0.583 | 0.052 |
